# Interactions of N- and C-terminal parts of Ana1 permitting centriole duplication but not elongation

**DOI:** 10.1101/2024.10.28.620588

**Authors:** Agota Nagy, Levente Kovacs, Helene Rangone, Jingyan Fu, Mark Ladinsky, David M. Glover

## Abstract

The conserved process of centriole duplication requires establishment of a Sas6-centred cartwheel initiated by Plk4’s phosphorylation of Ana1/STIL. Subsequently the centriole undergoes conversion to a centrosome requiring its radial expansion and elongation, mediated by a network requiring interactions between Cep135, Ana1/Cep295, and Asterless/Cep152. Here we show that mutant alleles encoding overlapping N- and C-terminal parts of Ana1 are capable of intragenic complementation to rescue radial expansion. This permits recruitment of Asl and thereby centriole duplication and mechanosensory cilia formation to restore the coordination defects of these mutants. This genetic combination also rescues centriole duplication in the male germ line but does not rescue the elongation of the triplet microtubule-containing centrioles of primary spermatocytes and consequently these males are coordinated but sterile. Such centriole elongation is rescued by the continuous, full-length Ana1 sequence. We define a region that when deleted within otherwise intact Ana1 does not permit primary spermatocyte centrioles to elongate but still allows recruitment of Asl. Our findings point to differing demands upon the physical organization of Ana1 for the distinct processes of radial expansion and elongation of centrioles.

**IMPACT STATEMENT:** The centriole can undergo radial development and duplication using separated parts of the conserved Ana1 protein whereas elongation of centriolar microtubule triplets requires the continuous Ana1 primary sequence.

## Introduction

Centrioles are conserved organelles required to form centrosomes, the microtubule organizing centers of interphase and dividing cells, and as basal bodies to form cilia, which have multiple roles in motility and signalling (Azimzadeh, 2014; Loncarek and Bettencourt-Dias, 2018). Their diverse functions are reflected in their association of their misfunction with many diseases; cancer cells are known to have multiple centrosomes that contribute to the generation of aneuploidy in mitosis, centrosome function goes awry in the microcephalies, and the many inherited ciliopathies reflect the widespread importance of cilia in signalling processes (Jaiswal and Singh, 2021; Qi and Zhou, 2021) .

In the great majority of cell types, centriole duplication is restricted to once per cell cycle. The duplication process is promoted by Polo-like kinase 4 (Plk4) (Bettencourt-Dias *et al*., 2005; Habedanck *et al*., 2005; Kleylein-Sohn *et al*., 2007; Peel *et al*., 2007; Rodrigues-Martins *et al*., 2007) which is recruited to the flank of the daughter centriole by binding to Asterless (Asl)/ Cep152 (*Drosophila/* human nomenclature throughout) (Dzhindzhev *et al*., 2010; Kim *et al*., 2013; Sonnen *et al*., 2013) Plk4 phosphorylates Ana2/STIL in two regions; in the N-terminal part to enable Ana2 to be recruited to the mother centriole, and in its conserved STAN motif to enable Ana2’s binding to, and so recruitment of, the major cartwheel component, Sas6 (Dzhindzhev *et al*., 2014, 2017; Ohta *et al*., 2014) Assembly of the centriolar microtubules at the periphery of stacked Sas6 cartwheels and centriole growth then requires Sas-4/CPAP (Pelletier *et al*., 2006; Hsu *et al*., 2008; Tang *et al*., 2011; Cottee *et al*., 2013; Hatzopoulos *et al*., 2013)

At the onset of mitosis, cells have two centrosomes, each composed of one mother and one daughter centriole. During mitosis, a conserved centriole protein known as Ana1 in *Drosophila* and Cep295 in human cells plays a central role in centriole to centrosome conversion. In this process, the daughter centriole matures to become capable of recruiting peri-centriolar material (PCM), a complex molecular condensate that can nucleate cytoplasmic microtubules, and initiating duplication in the subsequent G1 phase (Izquierdo *et al*., 2014). We have previously documented how, in Drosophila, the inner centriole protein, Cep135, has its C-terminal part at the centriole’s core and its N-terminal part extending outwards to interact with the N-terminal part of Ana1. Ana1 in turn extends further outward with its C-terminal part interacting with the C-terminal part of Asl. These three molecules are sequentially recruited to the daughter centriole such that, following its disengagement from the mother centriole, the daughter becomes capable of assembling PCM and recruiting Plk4, Ana2 and Sas6 to generate the next generation of daughter procentrioles in G1 (Fu *et al*., 2016). Ana1 recruitment also requires a complex formed between Ana3 and Rcd4 (Panda *et al*., 2020; Tian *et al*., 2021). The Rcd4 protein is also required for the cohesion of doublet and triplet microtubule blades although its precise molecular arrangement and mechanism of action within the centriole is not clear (Panda, Ladinsky and Glover, 2024).

*ana1* mutants display a lack of coordination as a result of failures in centriole duplication and subsequent formation of microtubule doublet-containing cilia in the mechanosensory chordotonal organs (Blachon *et al*., 2009; Saurya *et al*., 2016a). In addition to its function in centriole assembly, Ana1 also has a role in centriole elongation. Centriole length directly correlates with Ana1 expression, with reduced Ana1 levels leading to shorter centrioles, and Ana1 overexpression causing a centriole elongation phenotype (Saurya *et al*., 2016a) As part of this study, Saurya and colleagues also showed that expression of a GFP–Ana1 fusion lacking the N-terminal 639 amino acids can support centrosome assembly and cilium function and so rescue coordination of an *ana1^mecB^* mutant. This led them to conclude that the N-terminal part of Ana1 is not required for these processes. Here, we re-examine this finding in the light that *ana1^mecB^* is not a null mutant but has a premature termination codon allowing it to express the N-terminal part of Ana1 (Blachon *et al*., 2009). We then use a genetic approach to define an overlapping region between N- and C-terminal parts of Ana1 necessary for the fragments to drive centriole duplication and formation of mechanosensory cilia.

However, these fragments fail to permit elongation of the microtubule triplet-containing giant centrioles of primary spermatocytes. We further define a region that when deleted within otherwise intact Ana1 also fails to permit primary spermatocyte centrioles to elongate but still allows recruitment of Asl. We have therefore been able to define two distinct functions for Ana1; one involving the separate N- and C-domains required for the recruitment of Asl in centriole duplication, the other required for elongation without affecting the recruitment of Asl. Thus, distinct regions of Ana1 with differing physical organisation are required for PCM and Plk4 recruitment and for centriole elongation.

## Results

### Intra-genetic complementation of *ana1^mecB^*

We have previously described how Ana1 is recruited to the centriole through an interaction between its N-terminal part and the N-terminal part of the core centriole protein Cep135 that together with Asl form a linearly arranged trimer that extends from the core of the centriole to its periphery (Fu *et al*., 2016). In this study, we showed that an N-terminal fragment of amino-acids 1-935 could be recruited by Cep135. This led us to consider how this could relate to the subsequent conclusion of Saurya and colleagues (Saurya *et al*., 2016a) that the N-terminal 639 amino acids of Ana1 are not essential for centrosome assembly and cilium function reflecting the fact that expression of a C-terminal Ana1 fragment (amino-acids 640-1729) could rescue the coordination defects of the *ana1^mecB^* mutant. We noted, however, that, the *ana1^mecB^* mutant had been described to have a premature STOP codon in a position that could enable expression of the first 1129 amino acids of the Ana1 protein (Blachon *et al*., 2009). This led us to consider if the functionality of the N-terminal part of the protein is retained in *ana1^mecB^* and whether Saurya and colleagues had been observing intra-genic complementation.

To address the question of whether that the N-terminal part of the protein was truly needed for centriole duplication and formation of the neurosensory cilia, we generated a null allele of *ana1.* We did this using CRISPR/Cas9-directed mutagenesis to delete the region encoding the first 762 amino acids of the protein and replace it with a DsRed gene having a transcriptional terminator (Figure 1A). No transcription of *ana1* could be detected in these flies confirming that it is a null mutant (Figure 1-figure supplement 1A). In contrast to *ana1^mecB^* flies, which were all able to eclose from pupal cases and yet were still severely uncoordinated, the *ana1^null^* flies were so uncoordinated that only 41% of the flies could eclose from the pupal case (Figure 1C). Thus, some minimal function persists in *ana1^mecB^*, possibly representing low level readthrough of the stop codon, that enables flies to emerge from the pupal case but they are then too uncoordinated to properly walk, fly or mate (Figure 1-video 1). The eclosion and coordination phenotype of *ana^null^* was completely rescued by the introduction of a transgene expressing the full length Ana1 fragment (*endo-ana1*), tagged with RFP, under the control of the endogenous *ana1* promoter (Figure 1C, Figure 2 A,B).

**Figure 1.**
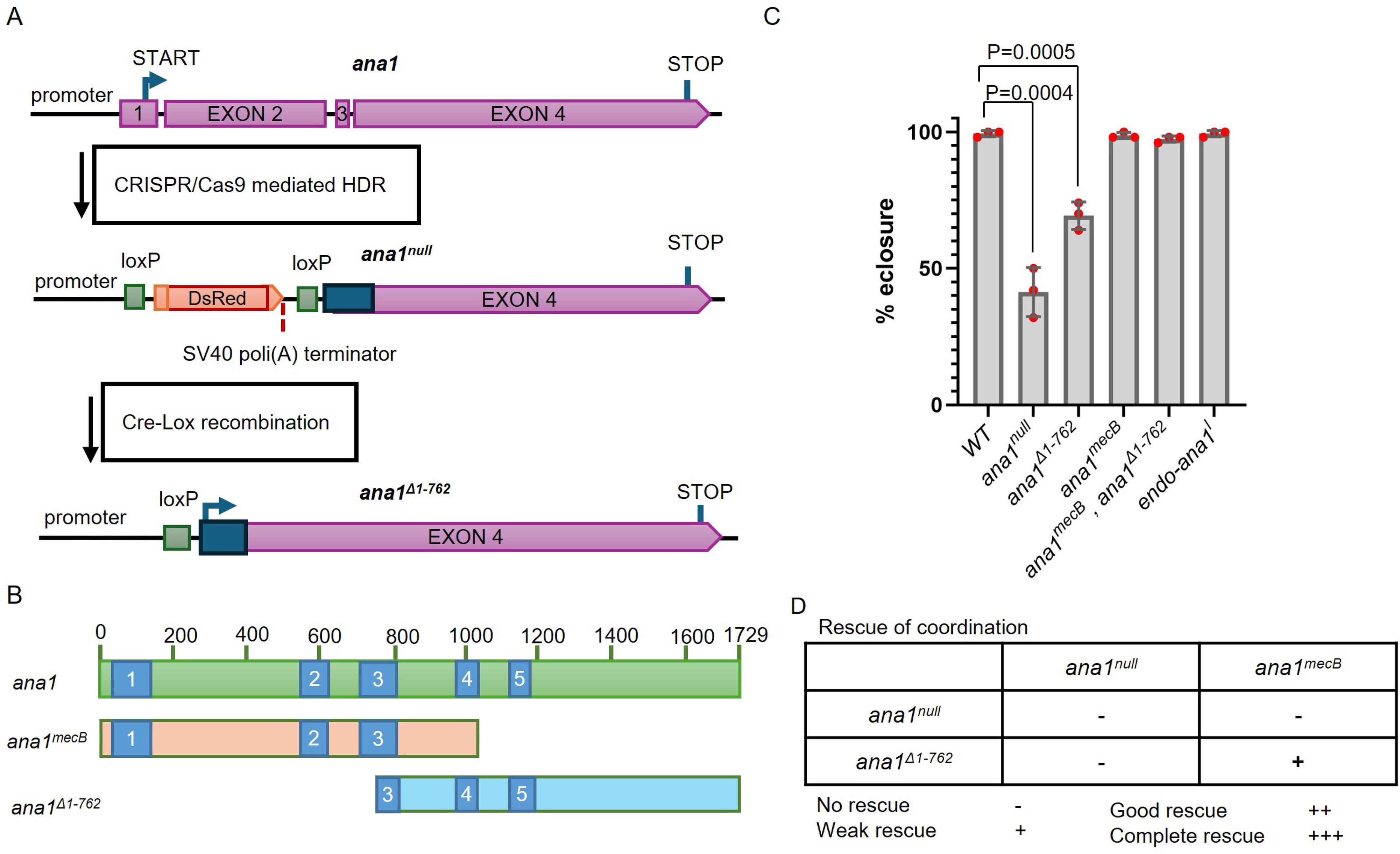
*ana1^mecB^* encodes an N-terminal Ana1 fragment, which can be complemented by an overlapping C-terminal fragment. **(A)** Schematic of *ana1* gene and mutant created by CRISPR/Cas9 mutagenesis. Exons 1, 2, 3 and a part of exon 4 of the gene were replaced with a *DsRed* marker, having a transcriptional terminator, using homology directed recombination (HDR). *loxP* sites flanking the DsRed marker permitted its removal via Cre-Lox-mediated recombination. This generates the *ana1^Δ1-763^* allele that expresses a C-terminal Ana1 fragment. **(B)** Schematic representation of Ana1 protein with five coiled-coil domains (blue) and the protein fragments encoded by *ana1^mecb^* and *ana1^Δ1-762^*. **(C)** Quantification of eclosion rate of indicated mutants. Pupae were aligned on the side of the vial and left to eclose, the number of adults were quantified. Flies were raised and tested at 25 °C. Means±SD are shown for n=150 flies of each genotype in N=3 independent replicates. *p* values of two-tailed, unpaired t-tests are shown for significant differences. **(D)** Complementation test for the coordination of the flies. Multiple tests and observations were made to assess the coordination of the flies, which were individually assessed under microscope without anaesthesia for each genotype. Cohorts of 10 flies were also transferred to an empty vial for climbing tests (Figure 1 supplement 1). All flies were raised and tested at 25 °C. No rescue (-), the flies couldn’t stand; weak rescue (+), the flies could walk, but not climb; good rescue (++), the flies can could with difficulty; complete rescue (+++), wild type phenotype.

**Figure 2.**
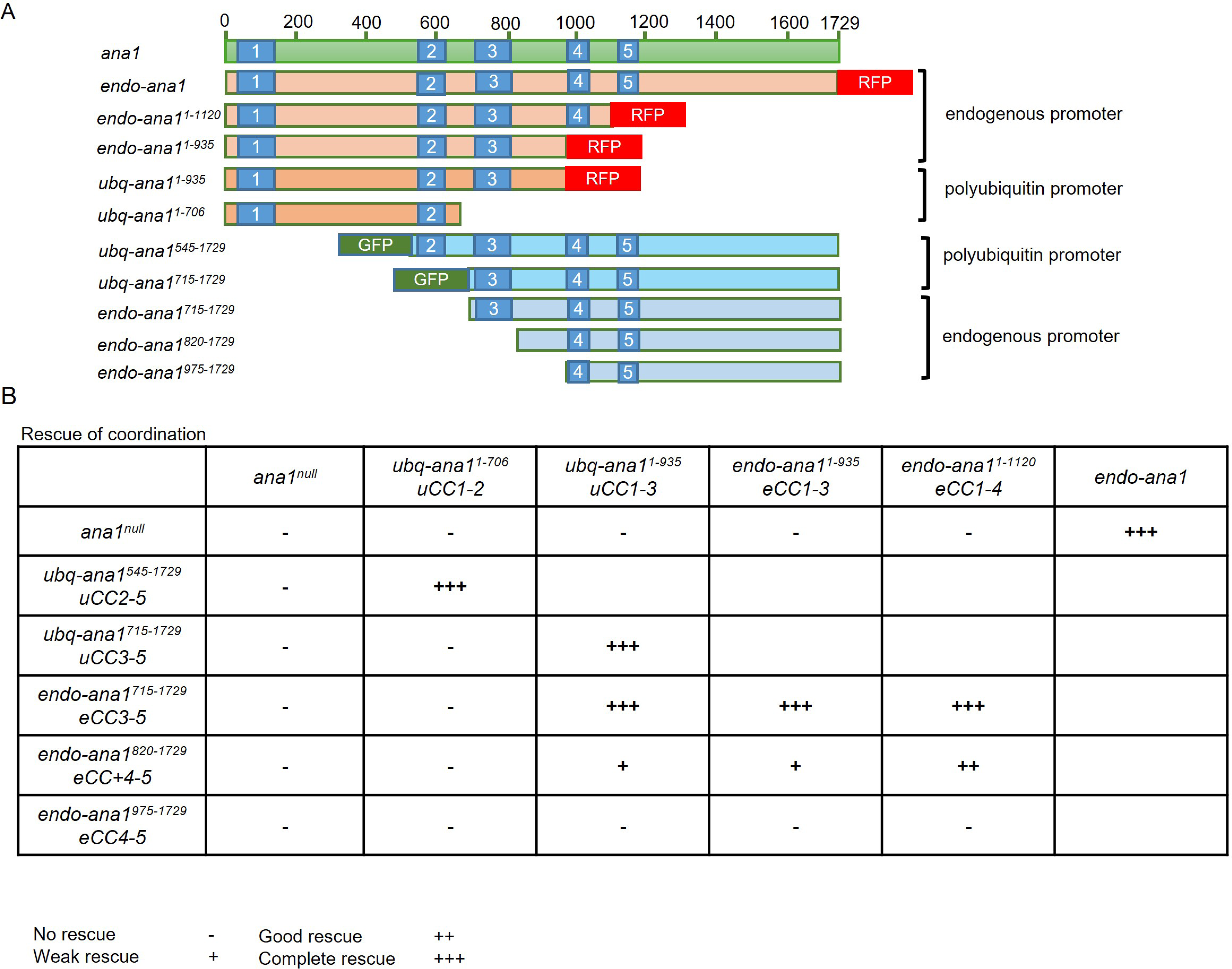
Expression of overlapping Ana1 N- and C-terminal fragments in an *ana1^null^* background can rescue coordination defects. **(A)** Schematic representation of N-and C-terminal fragments of different lengths under the regulation of the polyubiquitin promoter (ubq) or the endogenous ana1 promoter (endo). All fragments were expressed in an *ana1^null^* mutant background. Coiled coil domains (blue), GFP tag (green) and RFP tag (red) are indicated. **(B)** Coordination of flies expressing Ana1 fragments in different combinations. Individual flies of each genotype were observed under the microscope to assess coordination phenotype. Cohorts of 10 flies were also also examined in climbing assays. The number of individual flies that were able to climb 5 cm in 30 s were scored (Figure S2). Flies were raised and tested at 25 °C. No rescue (-), the flies couldn’t stand; weak rescue (+), the flies could walk, but not climb; good rescue (++), the flies can could with difficulty; complete rescue (+++), wild type phenotype. CC: coiled-coil domain; u: polyubiquitin promoter; endogenous promoter. The numbers following *CC* indicate the coiled-coil domains contained by the encoded Ana1 fragments.

Our design of the *ana1^null^* allele enabled removal of the transcription-blocking, DsRed cassette by Cre-recombinase mediated DNA recombination, enabling expression of a C-terminal part of *ana1* comprising amino-acids 763-1729 (*ana1^Δ1-762^,* Figure 1A,B). Around 69% of flies expressing this C-terminal part of Ana1 were able to eclose but once again, all were severely uncoordinated. However, when this C-terminal fragment was expressed in an *ana1^mecB^* mutant background, all flies eclosed and exhibited clear rescue, albeit partial, of their coordination (Figure 1BD, -figure supplement 1B; -video 1). This intragenic complementation of *ana1^mecB^* and *ana1^Δ1-762^* supports the hypothesis that *ana1^mecB^* directs the formation of an N-terminal fragment of Ana1 that when expressed with an overlapping C-terminal part is able to rescue Ana1’s function in coordinating movement.

The Ana1 protein contains five coiled-coil domains that have the potential to enable homodimerization of the molecule and so they also could potentially link the overlapping N- and C-terminal fragments of Ana1 studied here to provide a functional rod-like molecule.

Our *ana1^null^* mutant was designed so that Cre-lox mediated recombination would eliminate the DsRed gene to generate an *ana1^Δ1-762^*derivative that would specifically express the majority of exon 4 of the *ana1* gene. However, this strategy has the effect of interrupting coiled-coil domain 3 of the resulting C-terminal part (Figure 1B). We therefore considered the possibility that intact coiled-coil regions in this part of the molecule, including coiled-coil region 3, could be important to restore full coordination when independent N- and C-terminal parts were co-expressed. To test this hypothesis, we built a series of constructs encoding N-terminal parts of Ana1 of differing lengths that extended to include successive coiled-coil domains 2 to 4 (Figure 2A). We also built a set of complementary constructs of the C-terminal part extending N-terminally to include the same coiled-coil domain-containing regions. We introduced these into flies under the control of the endogenous *ana1* promoter using targeted φC31 integrase-mediated transgenesis to insert the transgenes encoding the N-terminal fragments into a defined region of the third chromosome and transgenes encoding the C-terminal fragments and the full length fragment into the second chromosome to ensure comparable levels of their expression. We also inserted some of these constructs downstream of the polubiquitin promoter to achieve higher expression levels. We then carried out a series of complementation tests to determine the extent of overlap between N- and C-terminal parts to give optimal rescue of coordination (Figure 2B, -figure supplement 1, -video-1). These intragenic complementation tests revealed that fragments overlapping in either or both the second and third coiled-coil domains plus the region immediately C-terminal of coiled-coil region 3 can completely rescue the coordination of flies (Figure 2B, -figure supplement 1, - video 2). We conclude that the presence of intact coiled-coil domains 2 or 3 enables the N- and C-terminal parts of Ana1 to interact and rescue Ana1’s function in providing coordination.

### Intragenic complementation of *ana1*’s N- and C-terminal coding regions re-establishes centriole duplication and mechanosensory cilia formation

Coordination in flies requires the function of type I ciliated mechanosensory neurons located in the femoral chordotonal organs (fChOs). Loss of coordination can reflect failure of the centriole duplication cycle to generate the basal bodies of these cells, failure of ciliogenesis, or both (Figure 3A). In the fChOs, neurons are clustered together in pairs, surrounded by an actin cone within structures known as scolopidia. Each neuron has a proximal basal body, derived from the daughter centriole, and a distal basal body, derived from the mother, which templates the cilia. We showed that fragments overlapping at coiled-coil domain 3 could completely rescue the uncoordinated phenotype. To test the extent of this rescue in the femoral chordotonal organs, we co-expressed fragments with the minimal overlapping sequences required to completely restore coordination. In control fChOs expressing full length Ana1 tagged with RFP, the proximal and distal basal bodies could be identified at the base of each cilium by both Ana1-RFP fluorescence and immunostaining to reveal dPLP (Figure 3B-lower row). In the severely uncoordinated *ana1^null^*flies expressing RFP-tagged Ana1^1-935^ having coiled-coil domains 1-3, we found irregularities in basal body numbers and the proximal and distal centrioles could not be distinguished, together indicative of a centriole duplication defect (Fig 3A - central). However, co-expression of RFP-tagged Ana1^1-935^ together with a 715-1729 aa C-terminal fragment, fully rescued coordination and restored the structure of the proximal and distal centrioles in the fChO. As these two Ana1 fragments overlap in coiled-coil region 3 and the sequences extending immediately C-terminal to it, it seems that these overlapping fragments can interact to restore a molecule capable of restoring the structure of basal bodies able to template cilia formation.

**Figure 3.**
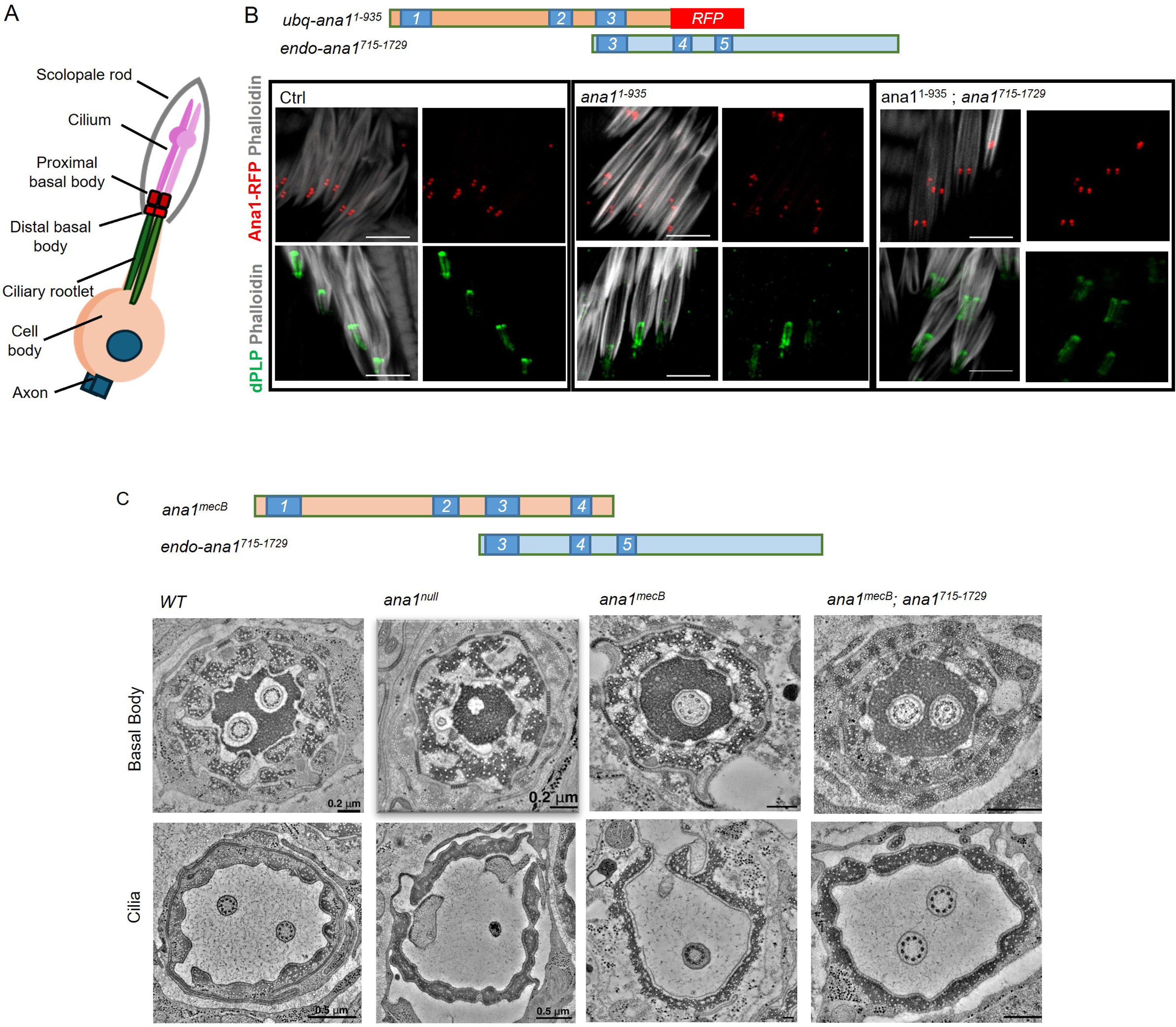
Complementary Ana1 fragments rescue basal bodies in the fChOs. **(A)** Schematic of a type one sensory neuron in *Drosophila* fCHOs. **(B)** Basal body organization in fChOs expressing full length Ana1-RFP (Ctrl), Ana1^1-935^ – RFP and Ana1^715-1729^ in an *ana1^null^* mutant background. The scolopidia were stained with phalloidin (white). Basal bodies were also marked with DPLP (green). n=5 fly legs from pharate adult flies were examined with similar results. Scale bar, 5 μm. **(C)** Transverse section of basal bodies and cilia in wild type (*WT*), *ana1^null^, ana1^mecB^* and *ana1^mecB^; ana1^715-1729^* femoral chordotonal organs imaged by electron microscopy.

To confirm the generality of the rescue of function by overlapping fragments and confirm its underlying cellular basis we used electron microscopy to examine the basal bodies and cilia of fChOs in more detail in *ana1^+^*, *ana1^null^*, *ana1^mecB^*, and *ana1^mecB^*; *ana1^C715-1729^* flies (Figure 3C, -figure supplement 1). *ana1^mecB^*; *ana1^C715-1729^* flies show a complete rescue of coordination. This is similar to the rescue seen in *ana^1-935^; ana^715-1729^* flies, reflecting the fact that both genetic combinations express the minimum overlapping sequence required to restore coordination. Transverse sections of scolopidia reveal them to contain a pair of basal bodies extending to a pair of ciliary axonemes in wild type fChOs as expected. The null mutant scolopidia showed a complete loss of basal bodies and cilia (Figure 3C, -video-1). On the other hand, we could find cases of *ana1^mecB^* mutant scolopidia that could have a remaining single basal body and cilium, some with normal structure (Figure 3C) and some with structural defects (Figure 3-figure supplement-1, -video-2). Expression of the C-terminal 715- 1729 amino acid fragment of Ana1 in the *ana1^mecB^* background fully rescued coordination and restored centriole and ciliary pairs in the fChO scolopidia (Figure 3C, -figure supplement 1). Thus, the C-terminally truncated Ana1 protein expressed in the *ana1^mecB^* mutant leads to defects in centriole duplication as it only expresses the first 1120 aa of the protein due to the premature STOP codon. However, the allele does retain a lower level of residual function in centriole duplication and cilium formation in the fChOs, that could either be due to this N-terminal fragment having some function *per* se, or to some readthrough of the stop codon. In either event, full function can be restored by expression of an overlapping C-terminal fragment.

### Interaction between the N- and C-terminal fragments rescues centriole duplication, but not elongation of triplet microtubules in the testes

The axonemes of *Drosophila*’s fChO cilia and the sperm develop from centrioles of quite distinct appearance. In the fChOs, for example, centrioles develop into basal bodies of approximately 120 nm in length from which the axonemes of the neurosensory cilia develop. Both basal bodies and cilia of the fChOs are built from doublet microtubules. In the male germline, a series of 4 rounds of stereotyped mitotic divisions with incomplete cytokinesis result in cysts of 16 cells having centrioles also of about 120 nm in length. The 16-cell cyst of spermatocytes undergoes an extended G2 phase of around 72 h during which time the two pairs of triplet microtubule-containing centrioles carried by each of these cells elongate to form so-called giant centrioles about 1.3 μm in length. The meiotic divisions segregate individual single centrioles into the spermatids which undergo elongation as the triplet microtubule containing axoneme develops into the sperm tail (Figure 4A).

**Figure 4.**
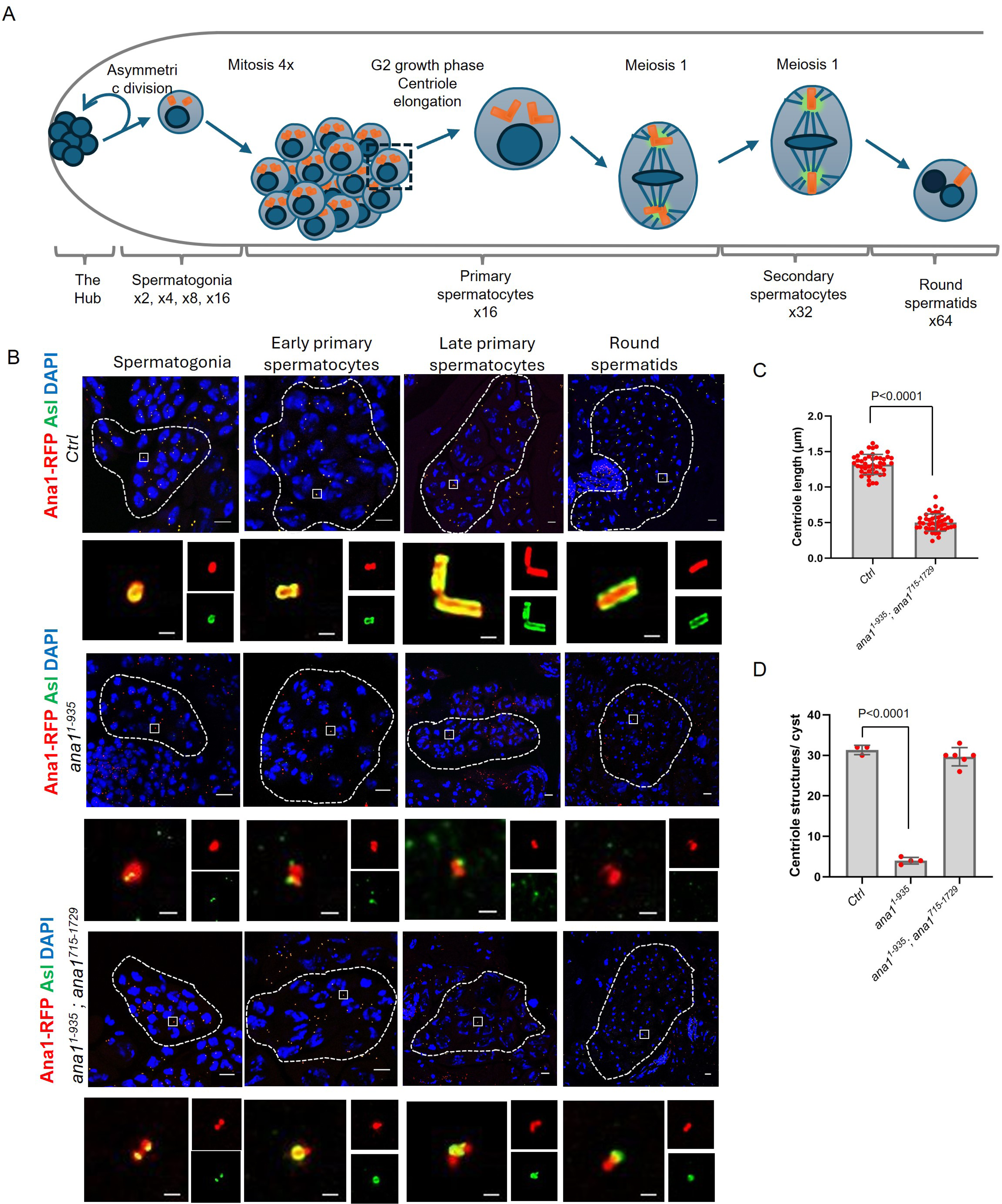
Complementary Ana1 fragments rescue Asl recruitment, but not centriole elongation in the testes. **(A)** Schematic of Drosophila spermatogenesis showing the progression of cells and their centrioles through the asymmetric stem cell division; the four rounds of mitosis that establish the spermatocyte cysts of 16 cells, each containing 2 centriole pairs; formation of the giant centrioles in the G2 growth phase of primary spermatocytes; the two meiotic divisions, which generate cysts of 64 spermatids, each containing a single centriole **(B)** Representative cysts of 16 cell spermatogonia, early and mature spermatocytes and the 64 cell cyst of round spermatids from *endo-ana1 (Ctrl)*, *ana1^1-935^* and *ana1^1-935^ ; ana1^715-1729^* flies. Centrioles are marked by Ana1-RFP or Ana1^1-935^-RFP (red) and Asl (green). Scale bar, 10 μm. Magnified examples of individual centrioles or paired centrioles are shown to represent the centriolar structures from each cyst. Ana1 or Ana1^1-935^ (RFP) and Asl (green) mark the centrioles. Scale bar, 0.5 μm. **(C)** Quantification of centriole length in *endo-ana1 (Ctrl)*, *ana1^1-935^; ana1^715-1729^* round spermatids using the Asl marker. n=45 centrioles were measured from N=3 testes. Means±SD are shown, *P* values of two tailed, unpaired t tests are shown for significant difference. **(D)** Quantification of centriole structures marked by both Ana1-RFP or Ana1^1-935^-RFP and Asl in *endo-ana1 (Ctrl)*, *ana1^1-935^* and *ana1^1-935^; ana1^715-1729^* mature primary spermatocyte cysts. n=3-6 cysts were scored from at least N=3 testes. Means±SD are shown, *P* values of two tailed, unpaired t tests are shown for significant difference.

We found that the co-expression of Ana1^1-935^ and Ana1^715-1729^ restored coordination to enable flies to mate but we were surprised to find that males of this genotype were sterile (Figure 4-figure supplement 1A). To determine the underlying cause of this sterility, we examined the formation of male germline centrioles and their behaviour in this *ana1^1-935^ ana1^715-1729^* strain in comparison to *endo-ana1*, and *ana1^1-193^* males (Fig. 4B). This revealed reduced centriole numbers in the spermatocytes (Figure 4D, -figure supplement 1C,2) of *ana1^1-193^*males that failed to elongate in the prolonged G2 phase of the primary spermatocytes. The overlapping N- and C-terminal fragments of Ana1 expressed in *ana1^1-935^; ana1^715-1729^* males rescued centriole duplication (Figure 4D, -figure supplement 1C) in the spermatogonia but the centrioles failed to elongate in primary spermatocytes (Figure 4B,C, -figure supplement 1-B). These short centrioles accumulated γ-tubulin and went through two rounds of meiosis (Figure 4-figure supplement 1D) to form round spermatids (Fig. 4B). Some of these spermatids underwent elongation and entered the seminal vesicle but they failed to mature into functional sperm thus accounting for the male sterility.

To test if we could rescue centriole elongation by extending the N-terminal part of Ana1, we expressed Ana1 N- terminal fragments comprising 1-1120, 1-1200 or 1-1430 amino-acids either alone or together with the 715-1729 amino-acid C-terminal fragment. Ana1^1-1120^ fragment contains 4 coiled-coil domains, and its length mimics the predicted protein expressed from *ana1^mecB^*, while both Ana1^1-1200^ and Ana1^1-1430^ contains all 5 coiled-coil domains. However, none of these fragments in any combination could rescue centriole elongation (Figure 4-figure supplement 1E). These results suggest that a full length Ana1 dimer, which could establish interactions between all of its coiled-coil regions, might be needed to establish the proper structure for elongation of these microtubule triplet-containing cilia.

### Requirement for a C-terminal part of Ana1 for centriole elongation

The failure of Ana1^1-1430^ to promote elongation in the above experiment suggested that the structural integrity of the last 300 amino acids of Ana1 is essential for this process. We argued that it might be possible to identify a region within the terminal 300 amino-acids that could provide this property. To this end we constructed transgenes able to express Ana1 deleted for amino-acids 1210-1340 (referred to as ΔAB); amino-acids 1341-1471 (referred to as ΔCD); and amino-acids 1472-1602 (referred to as ΔEF) and expressed these in an *ana1^null^* background (Figure 5A). We found that the centrioles in spermatocyte expressing Ana1-ΔAB did not differ significantly in length from control, wild-type centrioles. In contrast, we saw a 9% diminution of centriole length in spermatocytes expressing Ana1-ΔCD and a 54% diminution of centriole length following expression of Ana1-ΔEF (Figure 5B,D).

**Figure 5.**
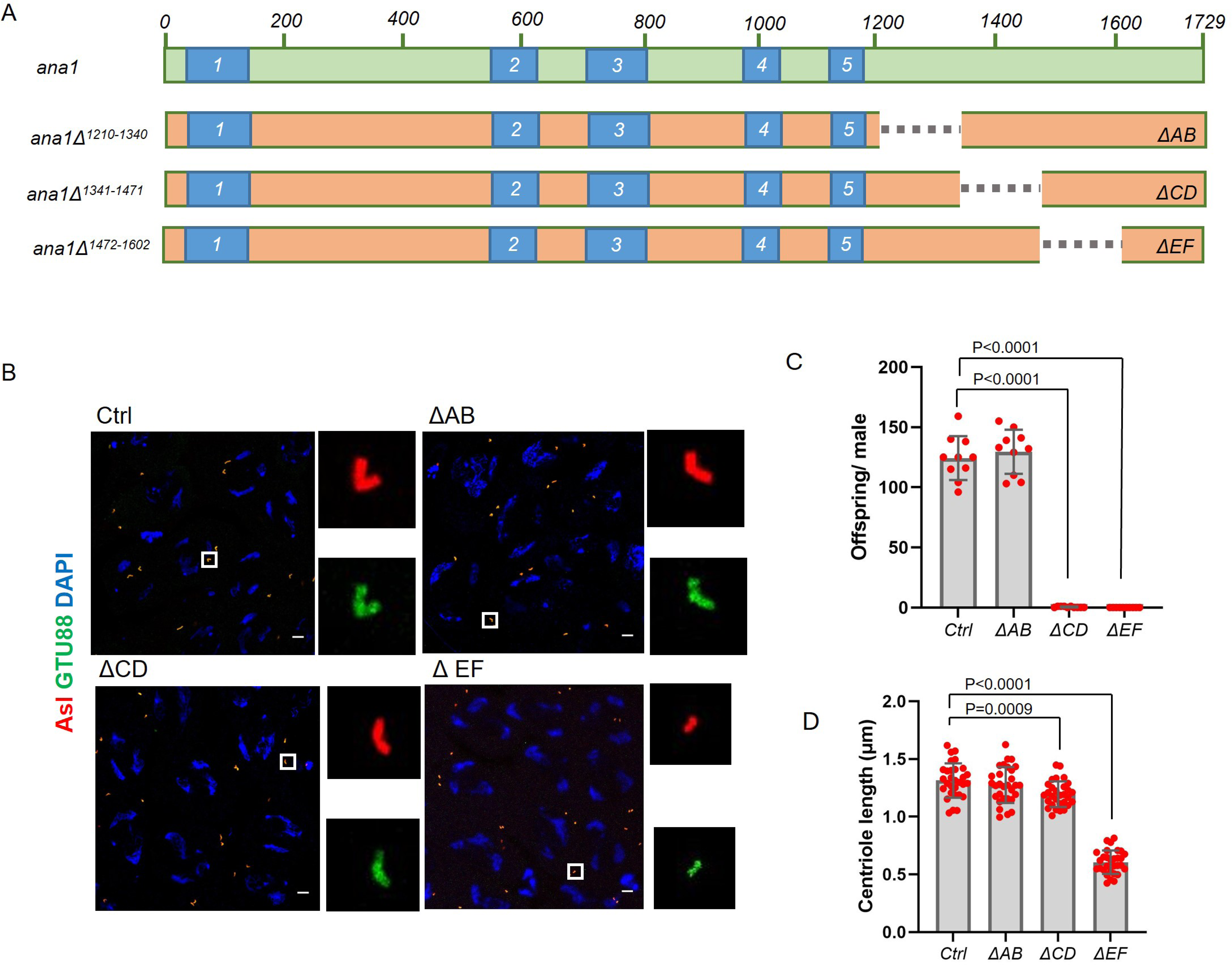
Sequential C-terminal deletions map the region of Ana1 essential for centriole elongation. **(A)** Scheme represents the site of deletions on Ana1 protein expressed from transgenes under^null^ the regulation of endogenous Ana1 promoter on ana1 mutant background. Deletions: *ΔAB,* deletion of aminoacids 1210-1340, *ΔCD*, deletion of aminoacids 1341-1471, *ΔEF,* deletion of aminoacids 1472-1602. **(B)** *WT* (Ctrl), ΔAB, ΔCD, ΔEF spermatocytes were immunostained to reveal Asl (Red) and γ-Tubulin, GTU88 clone (Green). Magnified examples of paired centrioles are shown to represent the centriolar structures from each genotype. Scale bar, 5 μm. **(C)** Fertility of wild-type (Ctrl), ΔAB, ΔCD, ΔEF males were tested by scoring the progeny of individual flies. Males were individually mated with wild-type females over 6 days in 25 °C. Means±SD are shown for n=10 flies per genotype, the *P* value of two tailed, unpaired *t* tests is indicated for significant difference. **(D)** Quantification of centriole length in *WT (Ctrl)*, ΔAB, ΔCD, ΔEF round spermatids using the Asl marker. n=30 centrioles were measured from N=3 testes. Means±SD are shown, *P* values of two tailed, unpaired t tests are shown for significant difference.

Interestingly flies expressing either Ana1- ΔCD or Ana1- ΔEF showed dramatic male sterility suggesting the possibility that the CD region might play a role in additional aspects of centriole function besides length control. That said, we conclude that Ana1 molecules that are fully capable of interacting with each other through the 5 coiled-coil domains require sequences in the EF region extending into the CD region to interact with other proteins required for the elongation of microtubule triplet-containing spermatocyte centrioles.

## Discussion

The term *centriole to centrosome conversion* was coined to describe how the daughter centriole is converted during mitosis into a structure able to nucleate cytoplasmic microtubules and duplicate, the centrosome (Wang *et al*., 2011). The process primarily requires recruitment of a rod-like array of molecules extending outward from the centriole’s core to provide a foundation on which to recruit *pericentriolar* material (PCM) to make a microtubule-organizing center (MTOC) (Wang *et al*., 2011; Fu *et al*., 2016). However, the concept of centriole to centrosome conversion has come to embrace centriole elongation because key centriole components, notably Drosophila Ana1 and its human counterpart, CEP295, are also required for centriole growth (Chang *et al*., 2016; Saurya *et al*., 2016a; Alvarez-Rodrigo *et al*., 2021). The genetic approach that we now describe predicts key aspects of the structure of Ana1/CEP295 that dictate the separable roles of this molecule in engaging Asl/Cep152 to recruit Plk4 and establish the PCM and in enabling the growth of centrioles that have triplet microtubules.

Our previous description of the assembly of the linear complex of Cep135, Ana1, and Asl of the *Drosophila* centriole provided a basis for centriole to centrosome conversion (Fu *et al*., 2016). Within this network, the N-terminal part of Ana1 physically interacts with the N-terminal part of Cep135, whereas the C-terminal part of Ana1 interacts with the C-terminal part of Asl (Figure 6). Our attention was therefore drawn to the conclusion of Saurya and colleagues that the N-terminal part of Ana1 is not required for centrosome assembly and cilium function in flies. This resulted from a finding that expression of a GFP–Ana1 fusion lacking the N-terminal 639 amino acids could support these processes in an *ana1^mecB^* mutant background (Saurya *et al*., 2016b). However, our present findings indicate that *ana1^mecB^* is not a null mutation. The original description of the *ana1^mecB^* allele would predict its ability to encode an N-terminal fragment of 1129 amino-acids (Blachon *et al*., 2009). Indeed, either *ana1^mecB^* or a DNA construct built to mimic this allele, but lacking the sequence 3’ to the stop codon, can be complemented through the independent expression of an overlapping C- terminal part of the molecule. Our findings with these and other constructs suggest that the optimal overlap between the N- and C-terminal parts to achieve this intragenic complementation include coiled-coil regions 2 and 3 and their immediate downstream sequences suggesting that this region is critical to achieve a stable overlapping interaction.

**Figure 6.**
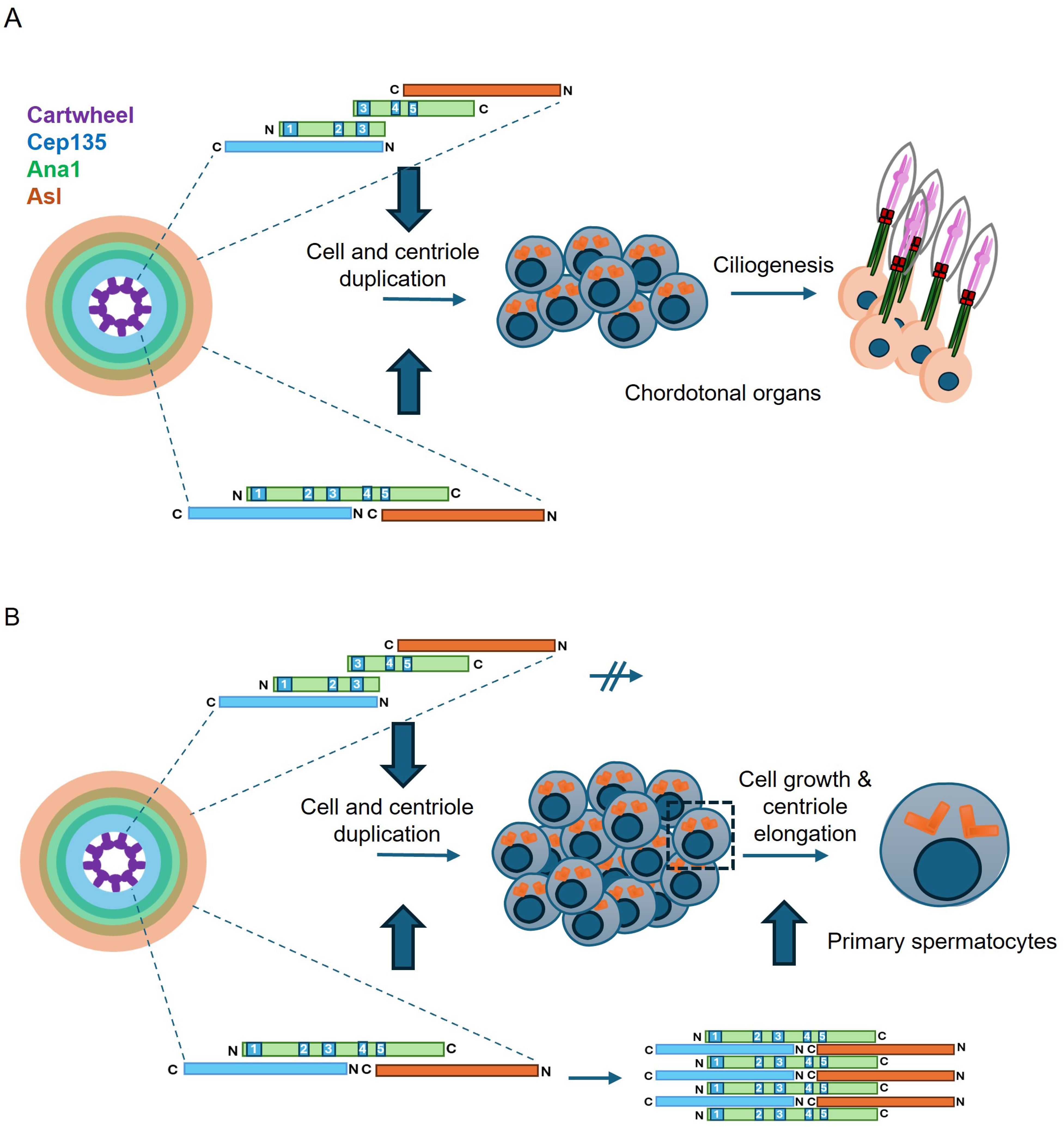
Model to account for differing physical organisaiton of Ana1 to permit centriole to centrosome conversion and elongation of centrioles with triplet microtubules. **(A)** Scheme shows overlapping N- and C-terminal parts of Ana1 interact through coiled-coil region 3 and flanking sequences to enable correct spatial recruitment of Asl as required for centriole to centrosome conversion, enabling centriole duplication and ciliogenesis. **(B)** The full length contiguous Ana1 sequence is able to interact between all coiled-coil regions giving the molecule potential to stack, thereby permitting elongation of triplet microtubules. Overlapping N- and C-terminal parts are structurally incapable of stacking and consequently do not permit centriole elongation. Elongation also requires Sas4 and Centrobin in a manner that remains to be determined.

This would establish linkage between four components - Cep135, the N-terminal Ana1 part, the C-terminal Ana1 part, and Asl (Figure 6) - permitting the correct spatial orientation of the Cep135-binding, N terminal part of Ana1 and the Asl-binding C-terminal part that are necessary for centriole to centrosome conversion. Moreover, the ability of the overlapping N- and C-terminal parts to restore formation of the sensory cilia of the chordotonal organs not only indicates rescue of centriole duplication, which requires centriole to centrosome conversion, but also, as we observed, rescue of the ability to assemble the chordotonal cilia that are built from doublet microtubules.

The independent expression of these overlapping N- and C-terminal parts of Ana1 was also sufficient to rescue the formation of 4 centrioles in the 16 immature primary spermatocytes. However, overlapping parts of Ana1 were insufficient to drive the elongation of centrioles in the extended G2 phase of primary spermatocytes of either *ana1^null^* or *ana1^mecB^* mutants.

Centriole growth in these mutants could, however, be rescued by full-length Ana1suggesting that the continuity of the Ana1 primary sequence is required for centriole elongation but not for centriole to centrosome conversion.

Whereas centrioles of somatic cells in *Drosophila* contain doublet microtubules, the male germ line exhibits centrioles and axonemes comprised of triplet microtubules. However, EM studies by two of us (HR and JF, data not shown) have shown that the short centrioles seen in *ana1^mecB^* mutant primary spermatocyte centrioles also displayed doublet microtubules. The expression of full length Ana1 not only rescues centriole elongation but also microtubule triplet formation. A number of other centriole proteins have been implicated as having functions in centriole elongation and in several of these cases, there is evidence of a link between centriole growth and formation of triplet microtubules. Overexpression of Sas4/CPAP, for example, results in elongated centrioles (Kohlmaier *et al*., 2009) whereas its deficiency results in centrioles with incomplete microtubule triplets that still can convert to centrosomes but fragment as the microtubule blades come apart (Vásquez-Limeta *et al*., 2022). Moreover, a specific mutation of CPAP has been described with reduced tubulin interaction that results in shorter centrioles and cilia with doublet and not triplet-microtubules (Zheng *et al*., 2016). The *Drosophila* Centrobin protein is also required to generate triplet microtubules and promote elongation of spermatocyte centrioles (Reina *et al*., 2018). Its mammalian counterpart, CNTROB is also not only required for centriole duplication, but also for elongation and microtubule stability through interactions with tubulin and CPAP (Jeong *et al*., 2007; Jeffery *et al*., 2010, 2013; Gudi *et al*., 2011, 2014, 2015; Shin *et al*., 2015). These requirements for Centrobin and Sas4/CPAP for centriole elongation and C-tubule stabilization to form microtubule triplets predict they will exhibit functional interactions with Ana1.

Indeed, Ana1 can interact indirectly with Sas4, which binds to Ana1’s partner, Asl (Dzhindzhev *et al*., 2010). However, future studies are required to investigate such possibilities.

Our findings lead to two hypotheses about the structural organisation of Ana1 that require testing in future structural studies. First, the continuity of the primary amino-acid sequence in a single polypeptide is not required for centriole to centrosome conversion *per se* because it can be achieved by separate N- and C-terminal fragments that overlap in the central coiled-coil region 3 and sequences C-terminal to it (Figure 6). This defines a minimal region required for the N- and C-terminal parts of Ana1 to interact and give full function in centriole duplication and establishment of the axonemes of sensory cilia with doublet microtubules. It suggests the possibility that the full length wild-type protein might form homodimers to achieve its normal function, although the most critical factor would be to achieve the correct spatial organisation of Cep135- and Asl-interacting regions as we have previously described (Fu *et al*., 2016). Second, our study points to a need for continuous Ana1 protein sequence throughout the region carrying the 5 coiled-coil domains to permit the coupled processes of doublet to triplet microtubule conversion and elongation of triplet-microtubule centrioles. We speculate that this might facilitate the longitudinal stacking of Ana1 and its complexed proteins to facilitate the elongation of triplet-containing microtubules of the giant spermatocyte centrioles (Figure 6).

Further studies are required to clarify the mechanism of both these processes. In both *Drosophila* and mammalian cells, the roles played by the Ana3:Rcd4 complex in recruiting Ana1/Cep295 remain poorly defined (Panda *et al*., 2020; Tian *et al*., 2021). Moreover, Atorino and colleagues (Atorino *et al*., 2020) have proposed that Cep295 plays a role in stibilizing the microtubule wall structure in human cells independently of Cep135, which together with the cartwheel is lost from centrioles in contrast to its retention in *Drosophila* cells. This suggestion would be in line with our hypothesis for a role of the stacked Ana1 ortholog complex in centriole elongation. Our studies with deletion mutants in the extreme C-terminal region identified a requirement for the region around amino-acids 1472-1602 (region EF) for centriole elongation. The extreme male sterility of not only flies expressing Ana1- ΔEF, but also those expressing Ana1- ΔCD suggests that the CD region might play a role in yet other aspects of centriole function, besides length control, that needs more investigation. Interestingly, the EF interval overlaps with a region identified as required to recruit Polo kinase to the distal part of the spermatocyte centriole. The mutation of 34 serine to threonine residues in potential Polo kinase binding sites in this region abolishes Polo recruitment and centriole elongation (Alvarez-Rodrigo *et al*., 2021). However, this study did not identify Polo’s substrates in the vicinity of Ana1. Thus, how Polo kinase might facilitate the potential interactions of Ana1 with itself, with proteins such as Centrobin and Sas4, or with other neighbouring centriole proteins to promote triplet microtubule formation and centriole elongation requires future investigation.

## Methods

### DNA constructs

Full length *Ana1* cDNAs and their fragments of varying lengths were used to generate entry clones using the Gateway System (Life Technologies). Expression constructs were made by using the following destination vectors: pUWR, pUGW and a modified form of pTUW. In this modified vector, the promoter sequence was substituted with a genomic sequence of approximately 2000 bp from the *ana1* promoter region. Sequences encoding N-terminal Ana1 fragments were fused with C-terminal RFP using the Gibson Assembly system (New England Biolabs) before cloned into the modified vector.

Two guide RNA sequences were cloned into pCFD5 vector (Addgene, #73914). Left and right homology arm sequences of approximately 1000 bp flanking the gRNA target sites were cloned into the pHD-DsRED-attP-w+ vector (Addgene, #80898) for CRISPR/Cas9-mediated homology-directed repair (HDR).

### Drosophila melanogaster stocks

Flies were maintained at 25 °C on standard yeast and cornmeal medium. The *ana1^mecB^*mutant was described previously (Avidor-Reiss et al. 2004) and the *ana1^null^* was generated by CRISPR/Cas9 genome engineering summarised in Figure 1. The *ana1^null^* flies referred to in experiments were hemizygotes of the genotype *ana1^null^* /*Df(3R)Exel7356*. Transgenic flies were created by φC31 integrase mediated transgenesis to ensure comparable levels of expression. Transgenes encoding the N-terminal fragments were introduced into the BDSC_8622 stock, where they were integrated into chromosome III, while transgenes encoding the C-terminal fragments and the full-length protein were introduced into stock attP40, integrated into chromosome II. Flies expressing endo-Sas6-GFP were used to mark the centriole core. Wild-type control flies were *w^1118^*.

### Semiquantitative reverse transcription PCR

Total mRNA was extracted from 10 freshly eclosed adult males of each genotype using TRI Reagent™ Solution (ThermoFisher Scientific). Total mRNA solutions were treated with DNase I (New England Biolabs) to remove any traces of genomic DNA contamination. A cDNA library was generated by reverse transcription using the RevertAid First Strand cDNA Synthesis Kit (ThermoFisher Scientific). Equal concentrations of cDNA-s were added to a PCR reaction performed under non saturating condition (32 cycles). Primers specific for the *RpL17A* housekeeping gene were used as loading control.

### Eclosion test

40 pupae per genotype were gently transferred into new vials. The number of adults were recorded upon eclosion and statistically analysed. The test was repeated 3 times, and the data analysed using a two-tailed, unpaired T-test.

### Coordination tests

Pupae were transferred into new vials placed on their sides, to prevent the uncoordinated flies from sticking into the media. In order to test flies with weak coordination rescue, individual flies were transferred into a petri dish and examined under the microscope. The time (in seconds per minute) that flies were able to stand up and walk. Climbing tests were also performed for cohorts of ten flies. Before the assay, flies were transferred into an empty testing vial without anaesthesia. The vials were illuminated from above. The flies were tapped down to the bottom of the vial and given 30 seconds to climb up. The number of flies that crossed the 5 cm mark were recorded. Two independent cohorts of 10 flies were tested per genotype.

### Fertility tests

1–2-day old males were collected and individually mated with three wild type virgin females. The crosses were kept for 6 days, then the adults were removed. Vials in which any of the adults died were excluded. The number of offspring were recorded and statistically analysed. 10 crosses were evaluated per genotype.

### Immunostaining and confocal microscopy

To image femoral chordotonal organs, flies were raised until the pharate adult stage. The pupal case was removed, and the flies were fixed with 4% formaldehyde for 30 minutes. After washing three times in PBST, the upper part of the femur was finely dissected by opening the cuticle.

Testes were dissected from pharate adult males in PBS and transferred to PBS containing 5% glycerol for 20 seconds and then transferred on a microscope slide. The samples were squashed with a cover slip, frozen in liquid nitrogen and fixed in chilled methanol for 10 minutes. The samples were rinsed with PBS and washed with PBST for 10 minutes then incubated overnight with the primary antibodies in a humid chamber. The slides were washed three times 10 minutes in PBST and incubated with the secondary antibodies for three hours. The samples were washed three times and mounted with Vectashield+DAPI.

The following primary antibodies and dyes were used: rabbit anti-Asl (1:10000, recognises the N-terminus of the protein) (Dzhindzhev *et al*., 2010), mouse anti-γ-Tubulin antibody, clone GTU-88 (1:500, Sigma-Aldrich, T5326), chicken anti-dPLP (1:500) (Rodrigues-Martins *et al*., 2007), Alexa Flor 647 phalloidin (1:100). Secondary antibodies were diluted 1:500: goat anti-rabbit Alexa Fluor 488, goat anti-rabbit Alexa Fluor 568, goat anti-mouse Alexa Fluor 488 (Invitrogen).

All microscopic preparations were imaged using a Leica Stellaris 8 Falcon laser scanning confocal microscope under a 63X oil objective. Images were processed with ImageJ (Fiji).

### Centriole length measurements

The length of the centrioles was measured using the line profile tool in ImageJ (Fiji). To avoid ambiguity of tilted centrioles, only centrioles oriented perpendicularly to the imaging axis were measured.

### Statistical analysis

Means, SD, sample size (n) and population size (N) are indicated in the figures and figure legends. Data was analysed with two-tailed, unpaired *t*-test by GraphPad Prism (version 10.2.0). A 95% confidence interval was applied.

### Electron microscopy

EM was carried out as previously described (Panda et al. 2024)

## Acknowledgements

We are grateful to Pallavi Panda, Paula Coelho, and Ines Baiao-Santos for comments on the manuscript and, together with all members of the Glover lab, for stimulating discussions.

DMG is grateful for past support in Cambridge from the Wellcome Trust and currently in Pasadena from the National Institute of Neurological Disorders and Stroke of the National Institutes of Health under award R01NS113930.

---

**Figure 1-figure supplement 1.**
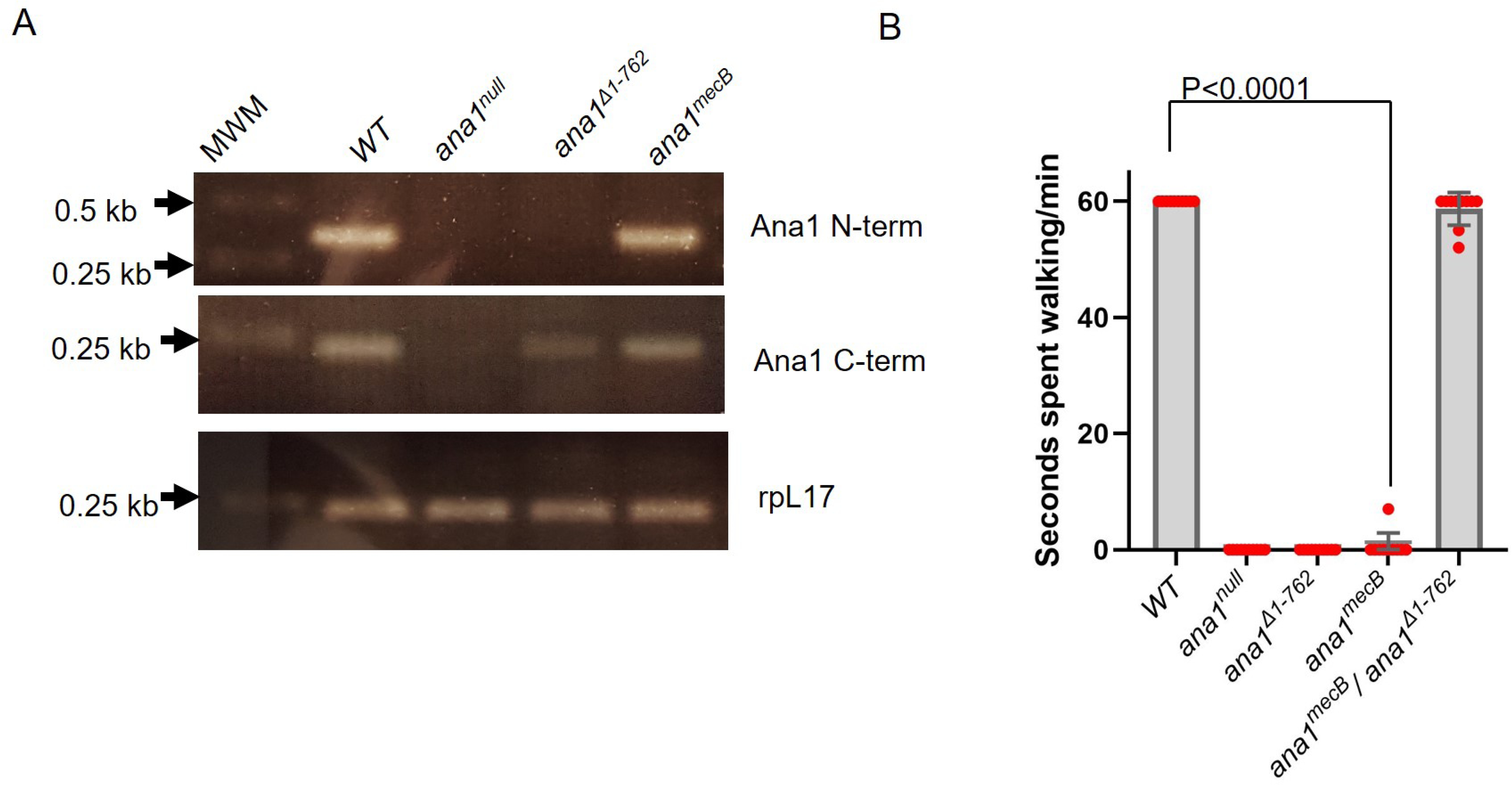
**(A)** Semiquantitative reverse transcription PCR shows transcription of the *ana1* gene in wild type (*WT), ana1^null^, ana1^Δ1-762^, ana1^mecB^* flies. Primers were specific to sequences encoding the N-terminal part and C-terminal part of Ana1. Total mRNA was extracted from freshly eclosed adult males for each genotype. cDNA library was created by reverse transcription. Equal concentrations of cDNA-s were added to a PCR reaction performed under non saturating conditions. Primers specific to *rpL17* were used for loading control. MWM: molecular weight marker; kb: kilobase. **(B)** Coordination assay for individual flies. The walking abilities of n=10 individual flies of each genotype were observed under microscope without anaesthesia. Flies were raised and tested at 25 °C. Means±SD and *p* values of two-tailed, unpaired t-tests are shown for significant differences are shown. For *WT vs ana1^null^* and *ana1^Δ1-762^ p* value can’t be calculated, as values in both columns are identical.

**Figure 2-figure supplement 1.**
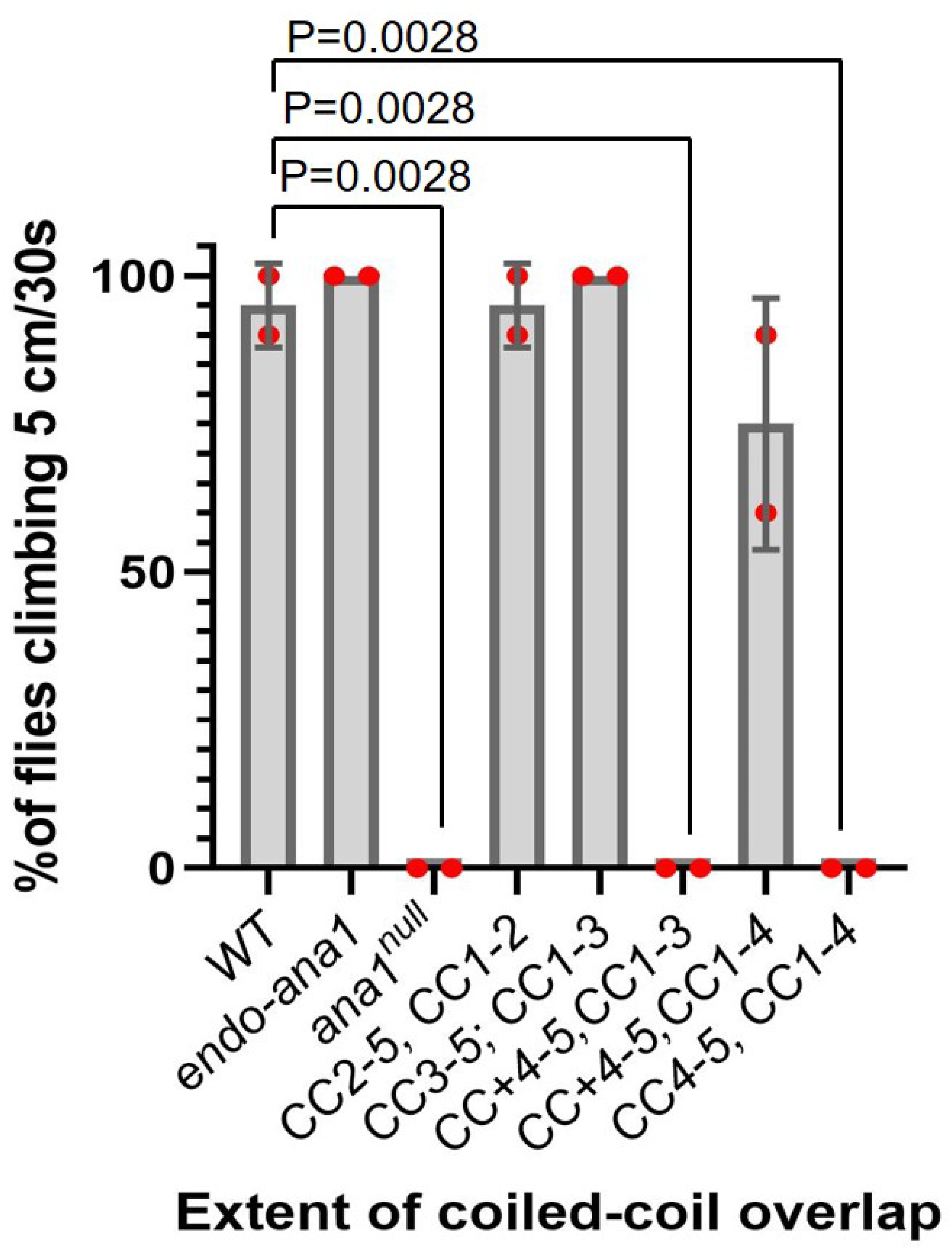
**(A)** Climbing assays of indicated transgenic flies expressing Ana1 fragments in an *ana1^null^* background. Cohorts of 10 flies were scored for their ability to climb 5 cm in 30 s. Means±SD are shown for n=10 flies were investigated in N=2 independent experiments. *P* values indicated for significant differences. Flies were raised and tested at 25 °C.

**Figure 3-figure supplement 1.**
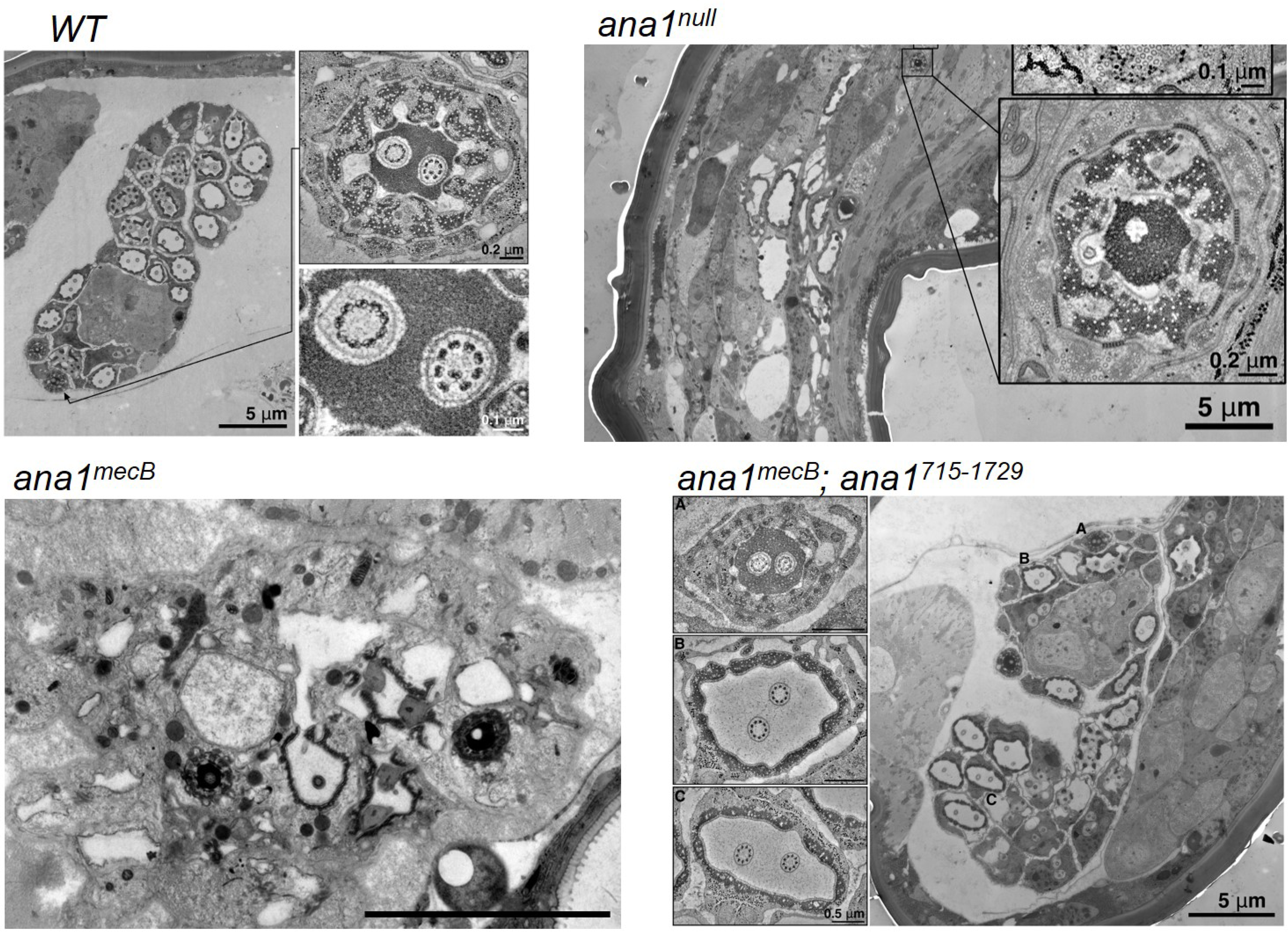
Overview of chordotonal organs in wild type (*WT*), *ana1^null^, ana1^mecB^* and *ana1^mecB^; ana1^715-1729^* femoral chordotonal organs imaged by electron microscopy.

**Figure 4-figure supplement 1.**
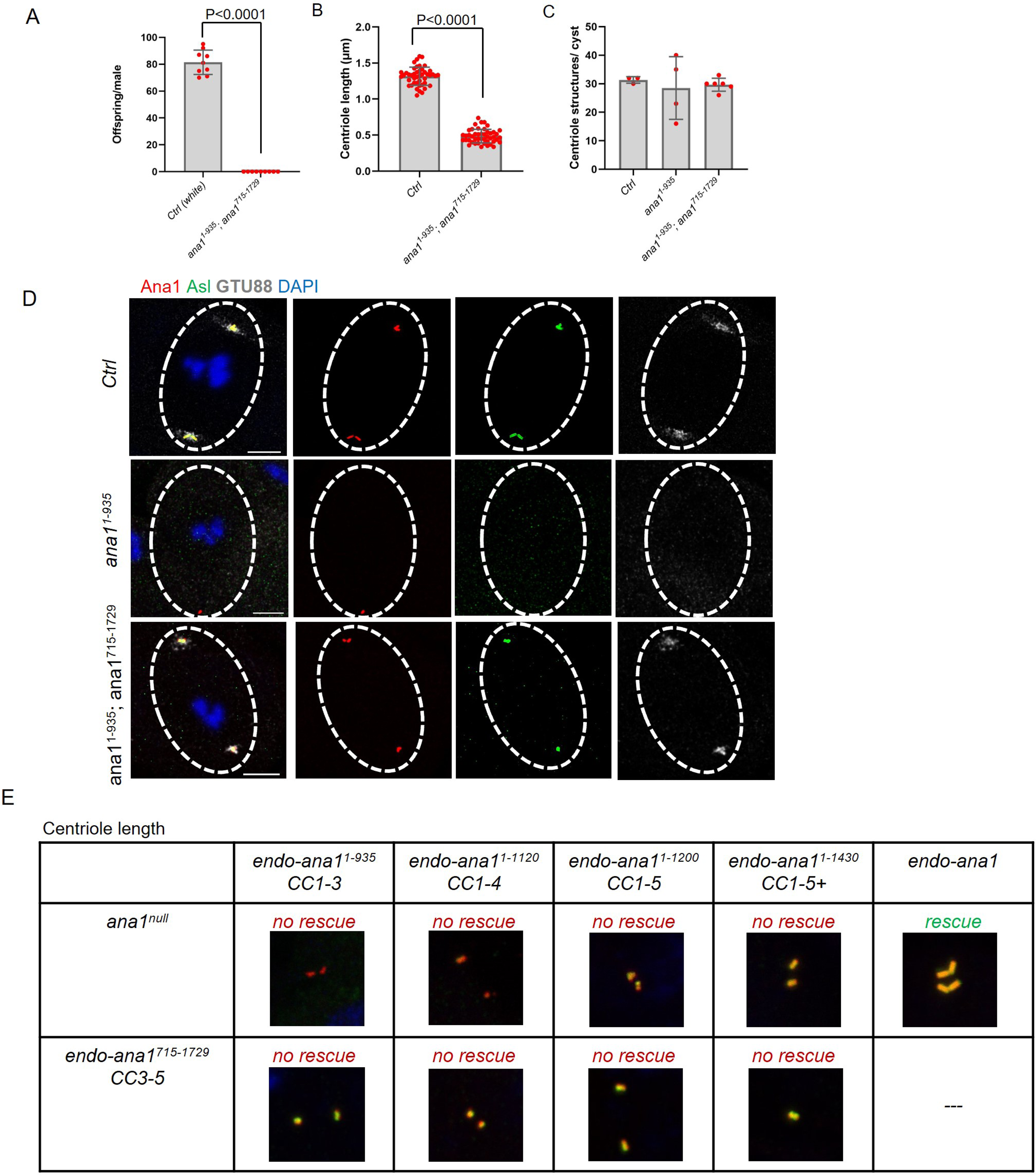
**(A)** Fertiliy of *WT* and *ana1^1-935^ ; ana1^715-1729^* males tested by scoring the progeny of individual flies. Males were individually mated with *WT* females over 5 days at 25 °C. Means±SD are shown for n=10 flies per genotype, the *P* value of two tailed, unpaired t tests is indicated for significant differences. **(B)** Quantification of centriole lengths in *endo-ana1 (Ctrl)*, *ana1^1-935^* and *ana1^1-935^; ana1^715-1729^* round spermatids using tAna1-RFP as marker. n=45 centrioles were measured from N=3 testes. Means±SD are shown, *P* values of two tailed, unpaired t tests are shown for significant differences. **(C)** Quantification of centriole structures marked by Ana1-RFP or Ana1^1-935^-RFP in *endo-ana1 (Ctrl)*, *ana1^1-935^* and *ana1^1-935^; ana1^715-1729^* mature primary spermatocyte cysts. N=3-6 cysts were scored. Means±SD are shown, *P* values of two tailed, unpaired t tests are shown for significant difference. **(D)** Representative primary spermatocytes in meiosis 1 from *endo-ana1 (Ctrl)*, *ana1^1-935^* and *ana1^1-935^; ana1^715-1729^* flies. n=3 meiotic cyts per genotype were examined from different testes with similar results. Centrososmes are marked by Ana1-RFP or Ana1^1-935^-RFP (red) and Asl (green) and γ-Tubulin, clone GTU88 (white). Scale bar, 10 μm. **(E)** Complementation tests scoring centriole length. Centrioles marked by Ana1-RFP (red) and Asl (green) were observed in primary mature spermatocytes. n=3 testes was examined per genotype with similar results.

**Figure 4-figure supplement 2.**
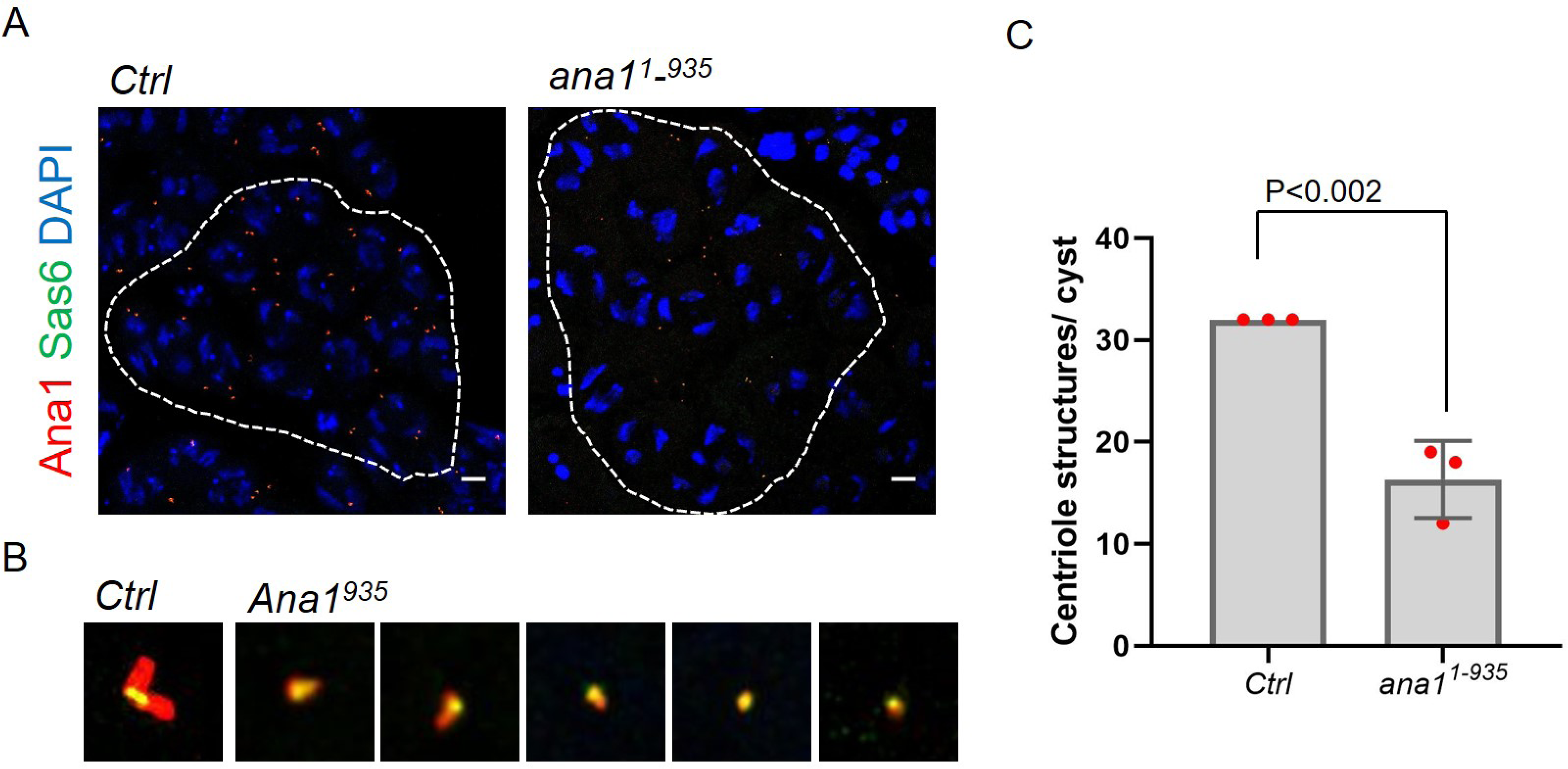
**(A)** Representative mature primary spermatocyte cysts expressing Sas6-GFP (green) together with full length Ana1-RFP (red) or the Ana1^1-935^-RFP fragment (red) in *ana1 an null* background. **(B)** Magnified examples of the centriole structures marked with Sas6 (green) and Ana1 or Ana1 1-935 fragment (red) **(C)** Quantification of centriole structures marked by both Ana1-RFP or Ana1^1-935^-RFP and Sas6-GFP in *endo-ana1 (Ctrl)*, *ana1^1-935^* mature primary spermatocyte cysts. n=3 cysts were scored from N=2 testes. Means±SD are shown, P values of two tailed, unpaired t tests are shown for significant differences.

**Figure 1-video 1** Coordination phenotypes of *ana1^mecB^, ana1^Δ1-762^* and *ana1^mecB^/ana1^Δ1-762^* flies. 6 flies from each genotype were transferred into vials without anaesthesia and their movements recorded. Flies were raised and tested at 25 °C.

**Figure 2- video 1** Representative videos for each coordination phenotype. 10 flies from each genotype were transferred to a petri dish and recorded. Flies were raised and tested at 25 °C. No rescue (-), the flies couldn’t stand; weak rescue (+), the flies could walk, but not climb; good rescue (++), the flies can could with difficulty; complete rescue (+++), wild type phenotype.

**Figure 2-video 2** Climbing assays of indicated transgenic flies expressing Ana1 fragments in an *ana1^null^* background. Cohorts of 10 flies were scored for their ability to climb 5 cm in 30 s. Flies were raised and tested at 25 °C.

**Figure 3-video 1** Movie shows Z stack volume of *ana1^null^* chordotonal organ transverse section revealing a missing basal body. Scale bar, 100 nm.

**Figure 3-video 2** Movie shows Z stack volume of *ana1^null^* chordotonal organ transverse section revealing cilia and a basal body. Scale bar 500, nm.

